# A deep learning approach to neurite prediction in high throughput fluorescence imaging

**DOI:** 10.1101/2021.04.23.441035

**Authors:** Mariya Barch, Melanie M. Cobb, Zachary Tokuno, Jen Leddy, Keili Prenton, Linus Manubens-Gil, Nicole Bellini, Stephanie Lam, Julia Kaye, Mara Dierssen, Steven Finkbeiner

## Abstract

Changes to neuronal morphology and loss of neurites and synaptic connections can be an important early indicator of neurological diseases, and a pathognomonic sign of neurodevelopmental disorders. These changes are typically detectable by microscopy in cell culture or histological samples, but quantification can be challenging. The neurites extending from cell soma can be quite thin, dim, overlapping and complex, making them laborious to trace manually and difficult to annotate and quantify computationally or automatically. Moreover, the tools available to aid this aim are limited in their capacity to generalize to high throughput image acquisition such as time-lapse or longitudinal imaging, where imaging conditions can change dramatically over the course of the experiment. In order to facilitate neurite quantification, we developed a deep learning (DL) neurite annotation prediction algorithm (NAPA) to predict the structure and length of neurites. NAPA overcomes experimental variation inherent to fluorescence imaging by learning more broader features that are important for neurite recognition. Based on a dataset with partial annotation, NAPA generated predictions on several unannotated datasets, and was able to capture differences between disease and control conditions. We also defined a sequence of steps to generate custom models with a small number of annotation inputs, and extended the predictions to a 3D tissue sample and longitudinal imaging. With this algorithm we developed an approach to quantify neurites with an accuracy that nears and sometimes exceeds human curation, in 1/100^th^ of the time. This approach makes accurate analysis of large or longitudinal datasets feasible across a broad range of datasets.

## Introduction

Neuronal morphology plays an important role in nervous system function, from how the brain forms connections in development to how the brain changes in response to disease. The number, dynamics and longevity of neuronal connections are informative to neuronal communication, synaptic plasticity, and normal brain function ^1^. The study of dendritic length and complexity, and spine density has become standard in the analysis of neuronal abnormalities, since alteration of these structures underlies many neurological diseases ^2^. Indeed, neuronal morphology can be a stronger marker than protein aggregates and plaques in determining early disease onset ^3–7^, and understanding the pace and extension of neurite loss could lead to novel therapeutic interventions for neurodegenerative disease ^8^.

In cell culture, neuronal soma size, neurite length, and branching complexity can all be measured ^9^ to evaluate physiological responses to perturbagens and potential treatments. However, accurately measuring neurite length and branching is challenging, as neurites are thin and irregular, and their dimness creates an inherent signal to noise constraint. These factors make hand tracing of neurites, the gold standard in neurite annotation, an extremely challenging task ^10^. This explains why neurites are not routinely assayed in studies and when they are, very limited sampling is performed.

To alleviate this manual annotation step and to improve the robustness and objectivity of the measurements, many groups have focused on automating some or all of the steps necessary for downstream neurite quantification ^11–13^. Available tools such as Imaris (Bitplane), FIJI Neurphology ^14^, and FIJI NeuronJ ^15^, and an implementation of a hidden Markov model ^16^ depend on image quality, culture complexity, and signal strength. However, human intervention is still required in all cases at least to ensure error checking and quality control ^17^.

Recently, deep learning (DL)-based techniques have been shown to learn fairly broad parameters directly from images, particularly with convolutional neural network (CNN)-based architectures. CNNs have been shown to recognize patterns in images that help classify objects and pixels effectively. Moreover, machine learning (ML) algorithms can reach human-level accuracy on specialized classification tasks ^18–20^. The feasibility of adapting natural image-derived networks to microscopy images has been shown for many classification tasks ^21–27^. This CNN-based approach can adapt to new contexts and requires fewer parameters, which the model learns to weight correctly on its own. CNN-based architectures may further weight parameters that human observers cannot perceive but are important in classifying microscopy images. In addition, manual tracing can help the model learn which pixels to ignore (e.g. out of focus objects, debris, or camera or plate artifacts) and which ones to include.

Here, we explore whether DL techniques can be used to automate manual annotation of neurites by learning relevant parameters across images with a broad range of imaging and experimental conditions. We describe a DL-based technique called NAPA (neurite annotation prediction algorithm) that takes a small number of annotated examples across a dataset and generates annotation for the full dataset. Since neurite tracing is a laborious manual process that can only capture neurites within the selected set of images, we designed a workflow that can expand and scale the investment of annotation to broader sets of images. This algorithm enables broad neurite prediction across large and diverse datasets and allows us to follow neurite connections longitudinally. In addition, by powering larger-scale neurite quantification, this approach can be used to probe the effect of potential therapeutics and address biological questions related to neurite structure in high throughput. NAPA enables neurite annotation with accuracies that approach and sometimes exceed hand curation and with a fraction of the time investment, making accurate neuronal structure analysis of large or longitudinal datasets feasible for the first time.

## Results

We set out to develop a model that learns the important elements in predicting neurites. Our approach was to build an auto-encoder shaped model, which has been previously used in applications such as image denoising ^28,29^. With this approach, we designate the neurite signal we want to learn, and all other sources of intensity in the images would be the noise we would like to remove (*e.g.* debris, soma, imaging artifacts). We prototyped this framework with available datasets that had already received varying degrees of neurite annotation (Table 1). It is worth noting that annotation tasks were not directly related to the modelling task presented here. In each case, we started with a set of images that had associated binary neurite trace masks. We then trained a model that could predict those masks. Finally, we quantified the accuracy of the predictions and the differences in neurite coverage across conditions. This sequence of steps is outlined in **Figure *1***.

**Table 1:** *The datasets used in this work and a summary of their parameters. All images are 16-bit depths, and table shows values derived from a histogram of one representative image*.

| Dataset | Cell type | Marker | Image<br>Min/Max<sup>5</sup> | Image<br>Mean±SD<sup>6</sup> | Annotation<br>Coverage<sup>7</sup> |
| --- | --- | --- | --- | --- | --- |
| I | DA<sup>8</sup> neurons<br>(human iPSCs) | AAV1- hSynapsin -<br>GFP | 0/13846 | 14±121 | Complete |
| II | Primary rodent<br>cortical neurons | pGW1-mApple | 24/3267 | 140±19 | Partial |
| III | CA1 pyramidal<br>neurons<sup>9</sup> | Fluorescent lucifer<br>yellow | 0/65535 | 6250±5516 | None/partial |
| IV | Forebrain neurons<br>(human iPSCs) | hSynapsin-mApple | 0/31400 | 27±252 | None/partial |

**Figure 1.**
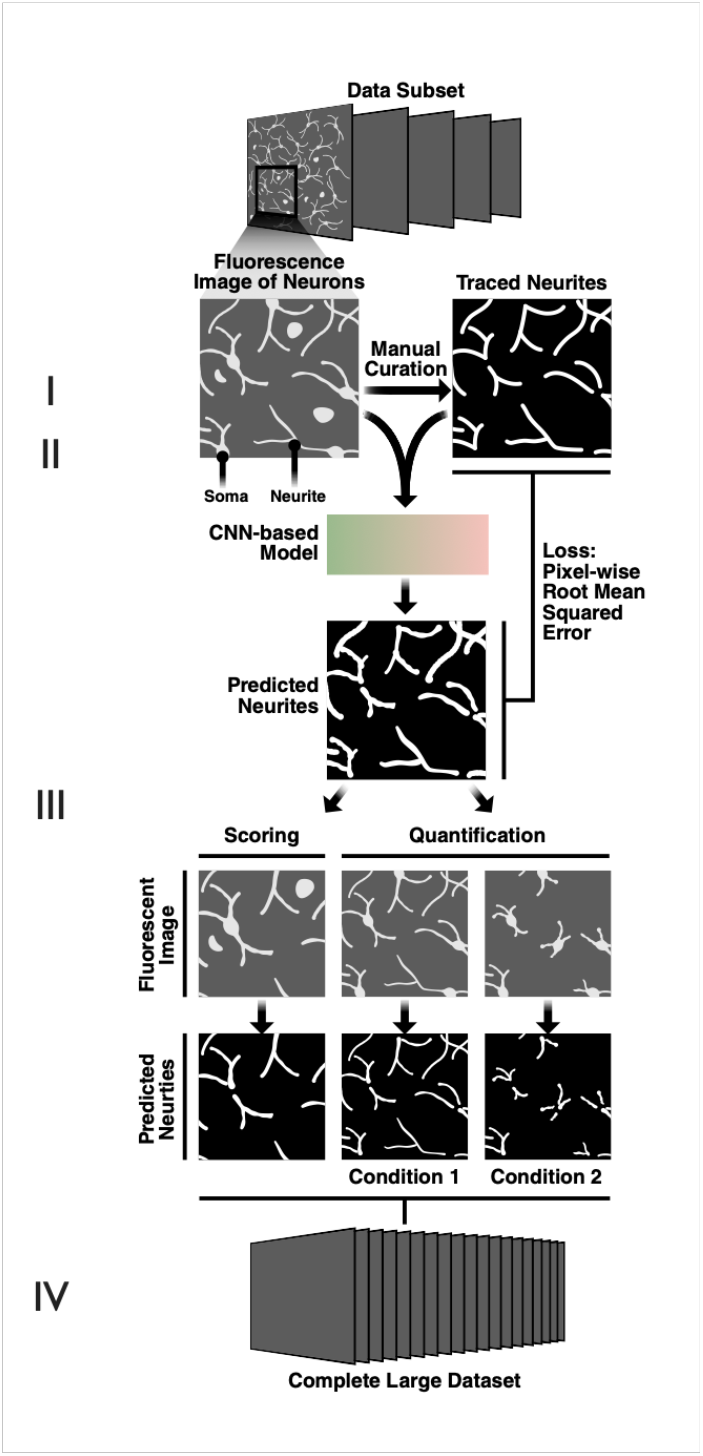
Schematic of workflow for generating NAPA. Images of cells in culture obtained using fluorescence microscopy are annotated through manual tracing of neurites, and these annotations that are converted to binary masks (I). Pairs of raw image intensities and corresponding binary masks (128 pixels x 128 pixels) serve as input pairs to NAPA (II). Pixel-wise root mean squared error (RMSE) loss is evaluated at each training step to minimize difference between prediction and target. Model output is used in downstream post-processing, scoring, and quantification steps (III) that can be applied to much larger datasets than the subset used for curation (IV).

The images in the datasets varied in image parameters (signal strength, amount of noise), sample preparation (cell type, cell species, method of whole cell marker expression, fluorophore and promoter used for cell marker, transfection efficiency, environment), imaging conditions (camera, microscope, exposure), and annotation coverage (full vs. partial). Representative images from a subset of the datasets are shown in **Figure *2*** and exemplify the variation we set out to capture.

**Figure 2.**
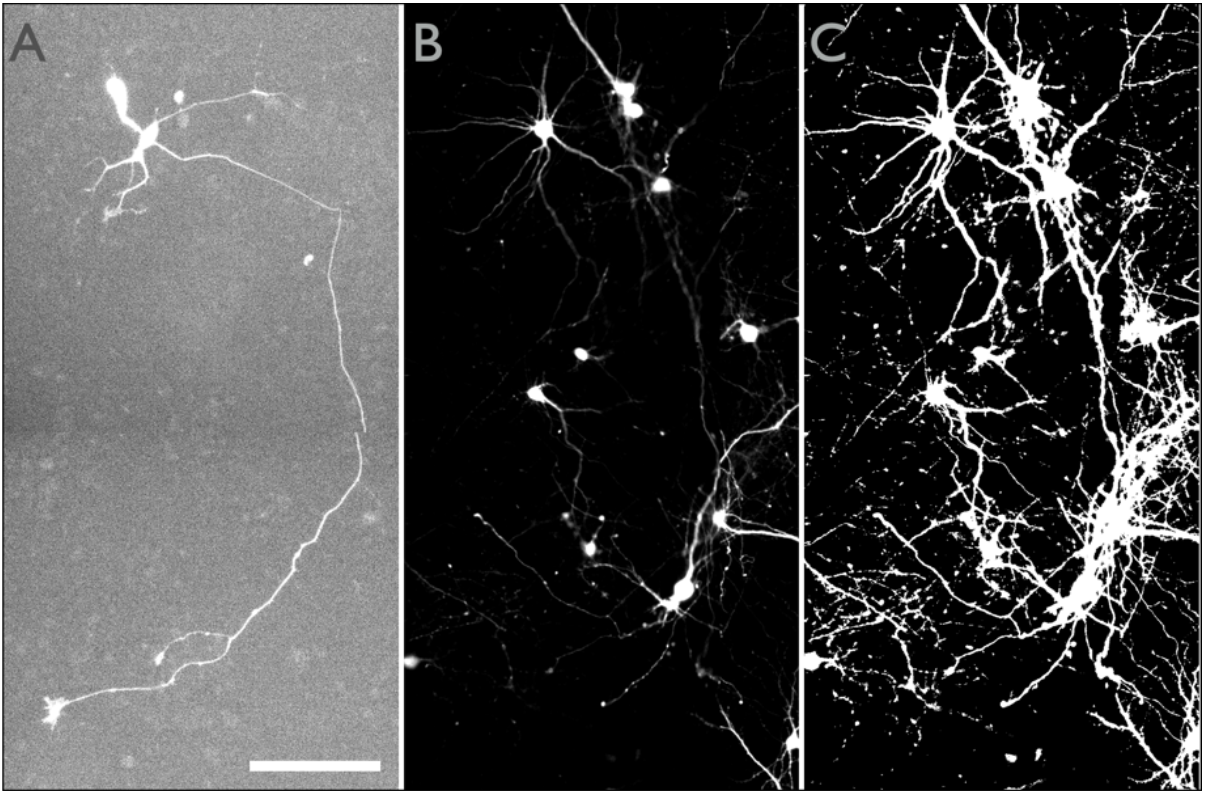
Example images of different neurons that demonstrate the variation in image acquisition across datasets. A. Neuron from dataset I that displays fairly low fluorescent signal. B. Multiple neurons from dataset II that display high fluorescent signal. C. Contrast saturated version of image in (B) enhancing the complexity of underlying neuronal morphology. Scale bar in A is 100um, scale is the same for all images.

We used a fully convolutional neural network architecture and explored model parameters that influence the ability to predict neurite annotation (**Figure *3***). Fully convolutional networks have been shown to work well in segmentation tasks for images of cells ^30^, so we asked if we could use a related architecture to selectively highlight the neurite structures in a fluorescence image. Pairs of images (the fluorescence image containing neurons and the corresponding traced neurite mask) served as input to predict an output that matched the annotation mask. The loss function, detailed in the methods, compares how closely the prediction and annotation match, per pixel, as an average root mean squared error (RMSE). The loss is further weighted to accommodate the imbalance of background (98%) and neurite (2%) pixels. Ultimately, the goal is to read in fluorescence images of cells that express the neuronal morphology marker and return masks that exclude soma and any debris, dust, or out-of-focus artifacts that might confound the fluorescence input.

**Figure 3.**
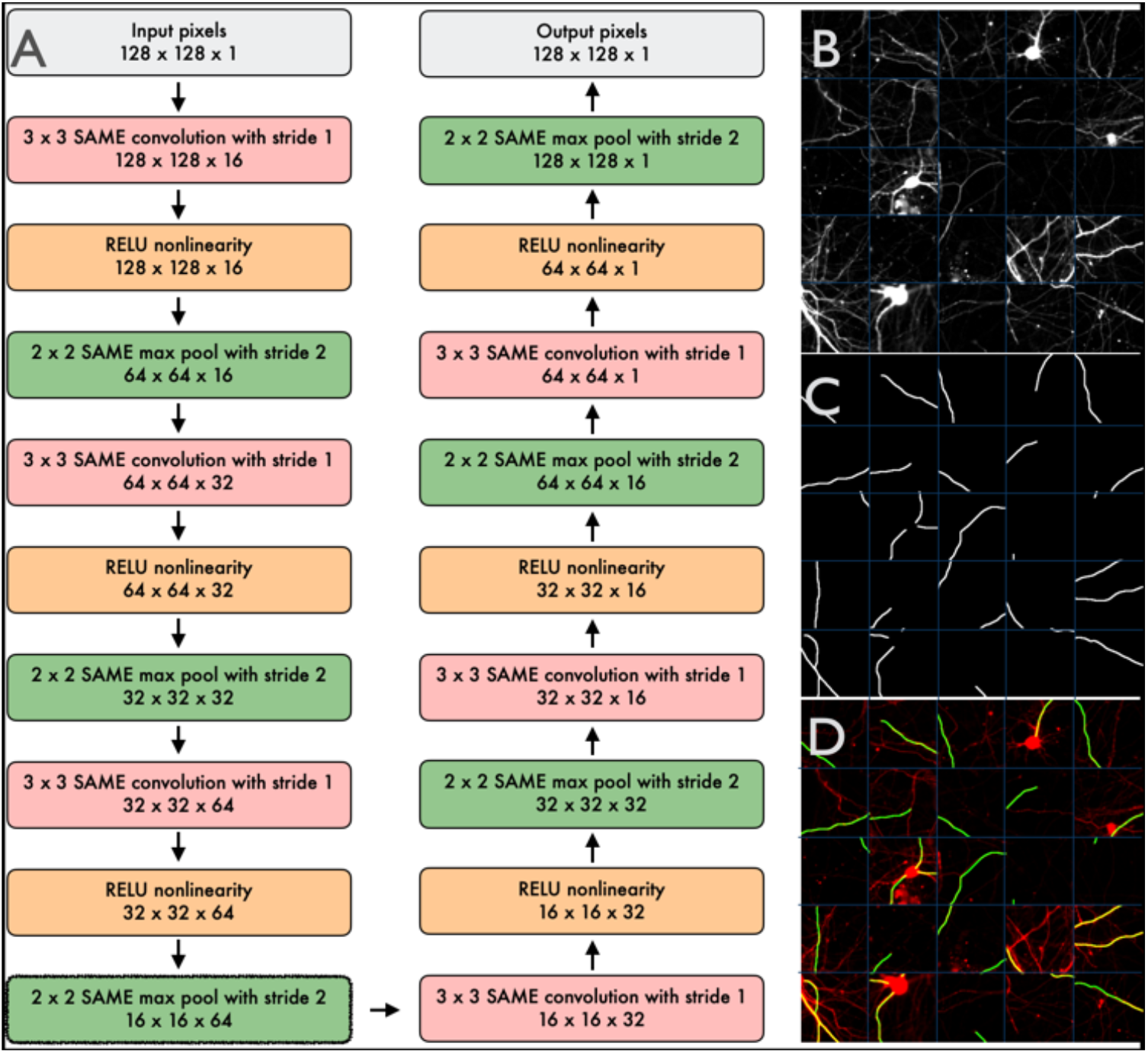
A. Schematic of model structure and example inputs. The CNN-based NAPA has three sets of convolution and pool layers in the contracting path, and three in the expanding path. Input is augmented with rotations, flips, and reflections. B–D. Examples of model inputs for a single dataset (dataset II). B. Twenty-five randomly chosen examples are used towards model generation. The 5 x 5 matrix shows patches of fluorescence images, with a faint grid added to visually separate image patches. C. Corresponding neurite traces converted to binary masks. D. Overlay showing partial coverage of the manually generated traces. Each tile in B–D is a 128×128 pixel patch and 40 μm wide.

To evaluate whether our model could learn to distinguish between neurite foreground and all-else background, we analyzed image patches with neuronal morphology marker and corresponding neurite annotation that were centered around soma (**Figure *4***A–C, dataset I). NAPA was able to learn to exclude soma and keep only neurites, and in some cases, it captured neurites better than manual curation. This dataset had sparse neurons, very low signal, and each neurite was annotated (complete annotation). We then extended this model to a partially annotated dataset (dataset II) of a denser culture with much stronger signal (**Figure *4***D–E). Notably, the model predicted unannotated neurites, indicating the potential of amplifying a given curation input and predicting the rest of the dataset. In addition, in the second dataset we used image patches that were no longer soma-centered, approaching a more realistic distribution of fluorescence in the full image. The encouraging result of predicting neurites from the more realistic input was indicative of the capacity to predict neurites in a full image.

**Figure 4.**
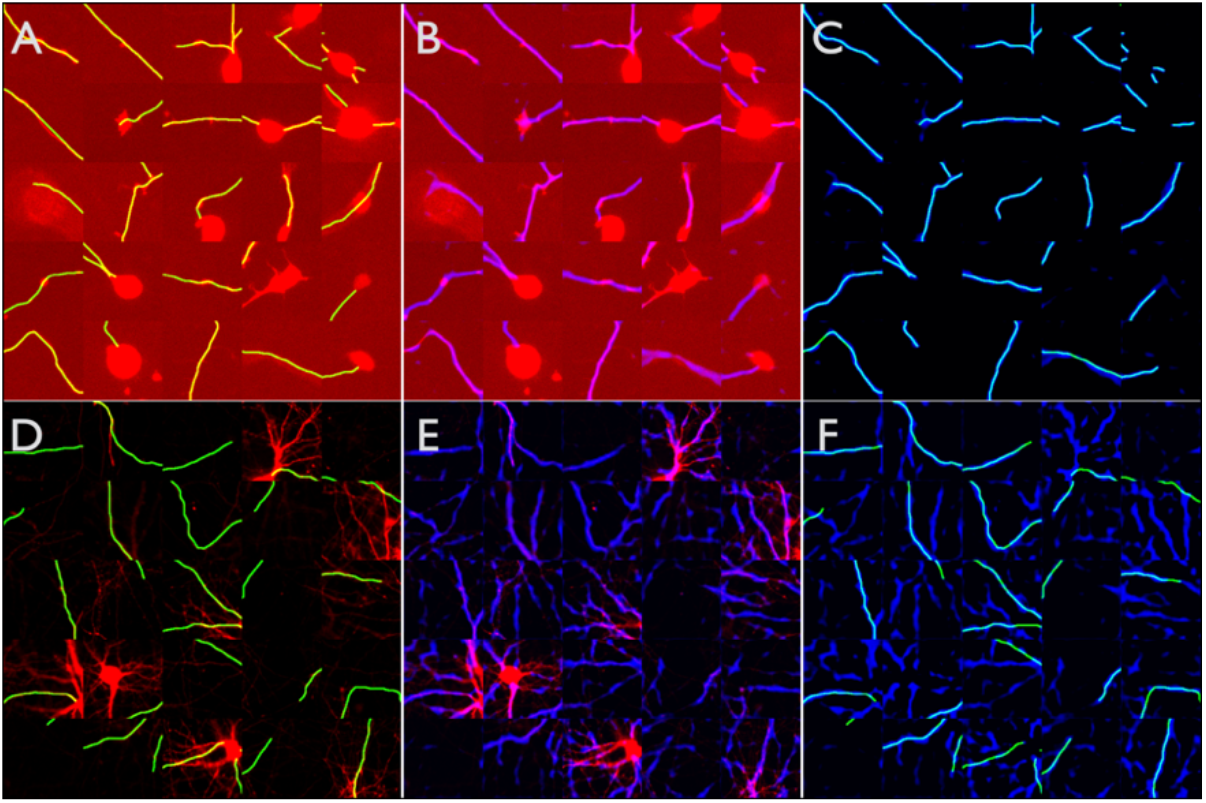
NAPA performs well on two prediction tasks from different datasets. A–C. Dataset I has low signal, fewer neurons, and every neurite is manually annotated. A 5 x 5 matrix of patches of fluorescence images (red) and manual annotation (green) overlays for dataset I (A), morphology images (red) overlayed with prediction (blue) (B), and prediction (blue) overlay with annotation (green) (C). D–E. In dataset II, the dominant neurites are annotated but many dim neurites are unannotated (partial annotation). A 5 x 5 matrix of patches of morphology images (red) and annotation (green) overlays (D), morphology images (red) overlaid with prediction (blue) (E), and prediction (blue) overlay with annotation (green) (F). The overlap between panel prediction and annotation is used to evaluate accuracy (Figure 7). Each tile in A–F is 40 μm.

Both for training and inference, we work with small image patches to streamline computation time. However, in full image prediction, patches can have more variation in content, ranging from no intensity, to bright spots of debris or soma, to complex networks of neurites, to everything in between. Therefore, we next asked if the model would continue to correctly assign neurites as foreground and everything else as background when considering full images. **Figure *5*** summarizes the result, where panels A and B show the full fluorescence and annotated images, respectively, and the predicted image is in panel C. An overlay of A–C is shown in panel **Figure *5***D, and a second example in **Figure *5***E. Neurite prediction for the complete image is computationally lightweight, and is much faster than manual annotation which can take several hours for images from a few columns of a 96 well-plate for a sparsely labelled sample. We applied this model to multiple other datasets with similar results (another example in **Figure *8***). Notably, as we tried crossing models trained on one dataset to predict on another dataset, we found that the best performance came from a model that was trained on the highest signal data.

**Figure 5.**
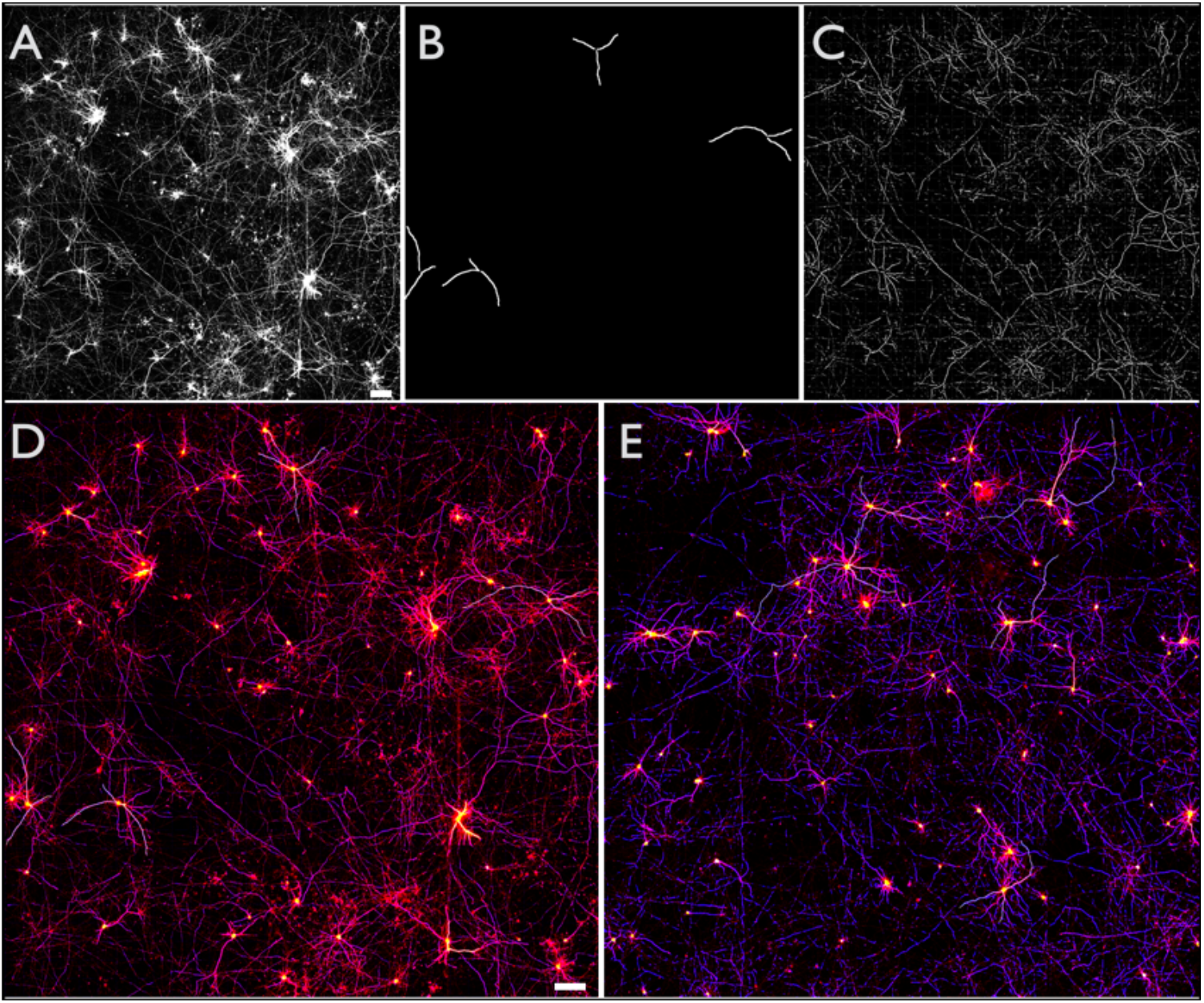
Neurite prediction for a full input image. Model is trained on small input tiles (128 x 128 pixels). Full image predictions shown in A–D. A. Fluorescence image of neuronal morphology marker. B. Corresponding annotation. For four cells, three of the most prominent neurites are traced. C. Output from model trained to predict neurite annotation. D. Colored overlay showing prediction (blue), morphology image (red), soma (yellow), and annotation ground truth (grey) for images A–C. E. Another image showing overlay with the same coloring scheme but enhanced prediction rather than fluorescence. Since the neurites are fairly thin structures, the images in are enhanced in the full view to give a sense of overall coverage. Scale bars in A and D are 100 μm.

**Figure 8.**
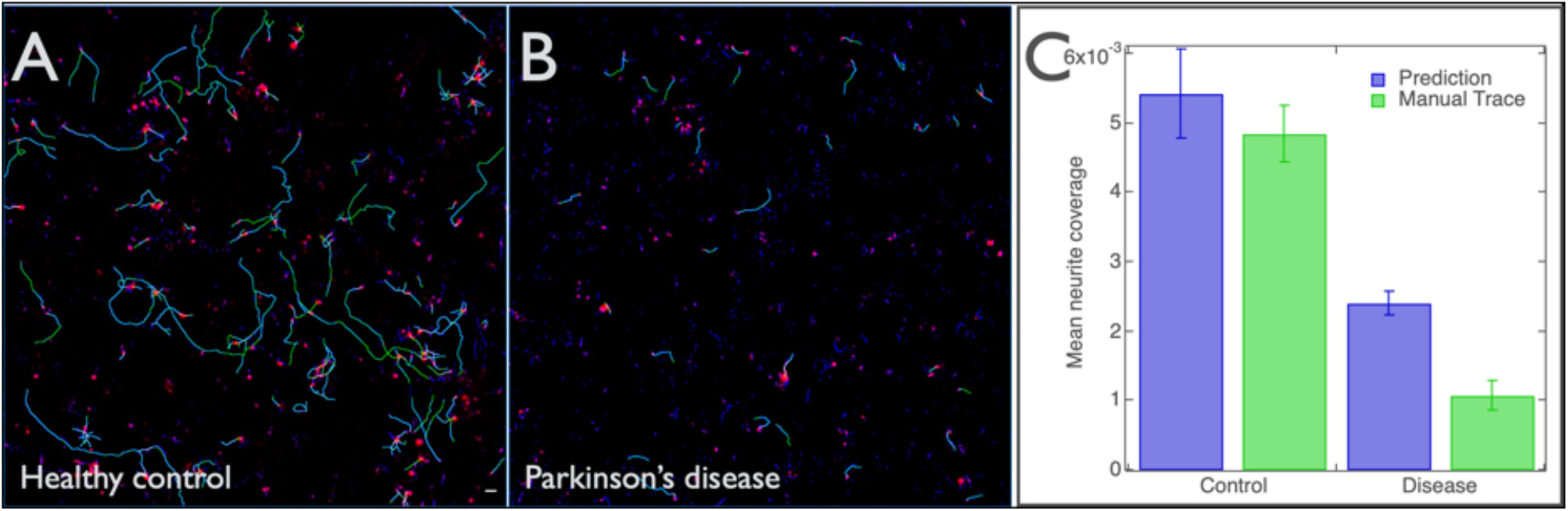
Differences between control and disease samples are evident by both manual and predicted neurite traces. Predictions were generated across 11 wells and used towards quantification of neurite coverage. A–B. Representative image overlays for iPSC-neurons derived from a healthy control (A) and Parkinson’s disease patient (B). Fluorescence image (red), manually traced neurites (green), predicted neurites (blue). Neurite contrast is artificially enhanced for visibility. C. Quantification of total neurite coverage (neurite pixels/all image pixels) for predicted and manually traced images for all images across disease (n=5) and control (n=6) groups. Scale bar represents 100 μm.

With these encouraging results across internal experiments, we next applied NAPA to neurons in a 3D tissue sample (dataset III). This dataset differs from the preceding data in several ways: the imaged neurons are from histology preparations of coronal brain slices, the tissue is imaged via confocal microscopy in a different laboratory, the image acquisition is low throughput and sampled in z rather than in the most focused x, y plane, and the sample is imaged through a glass coverslip rather than plastic multi-well plates. Our initial NAPA models did not extend well to this new dataset (data not included), which is perhaps not surprising, because our training data and its augmentations did not span the range of variation introduced by the new dataset. To develop a toolset that is more broadly useful, we sought to define a compact set of steps (or workflow) needed to generate data-specific models that could amplify training data by producing annotation for a complete dataset when curation of that scale is not generally possible.

One important requirement for such a workflow is that the amount of curation needed should be fairly minimal. We set out to determine the smallest number of curation examples that are necessary to produce a model that can infer the rest of the dataset. Initially, we tested this compact set of steps on dataset III, because it lacked annotation and thus we had no existing model for this dataset. In dataset III, there are only a few neurons in each image, and we tested tracing of 1, 2, 3, and 4 neurites per image for 20 images subsampled from 80 from a z-stack. Since this was our first purpose-annotated task, we defined criteria for what should be captured within the curation to minimize the number of training examples necessary to train a model that can label the remaining dataset. We annotated neurites that represented both dim and bright projection that clearly fell into focus within the image, and estimated that about 35 neurites (~400 128×128 training tiles) yielded good models. An example of prediction using a model for histology data from dataset III is shown in Figure 6, and a movie through the z-stack of the tissue is included as **Supplementary Movie *1***. We subsequently applied the same sequence of steps to other datasets from the lab, using NAPA to build individual models for each.

**Figure 6.**
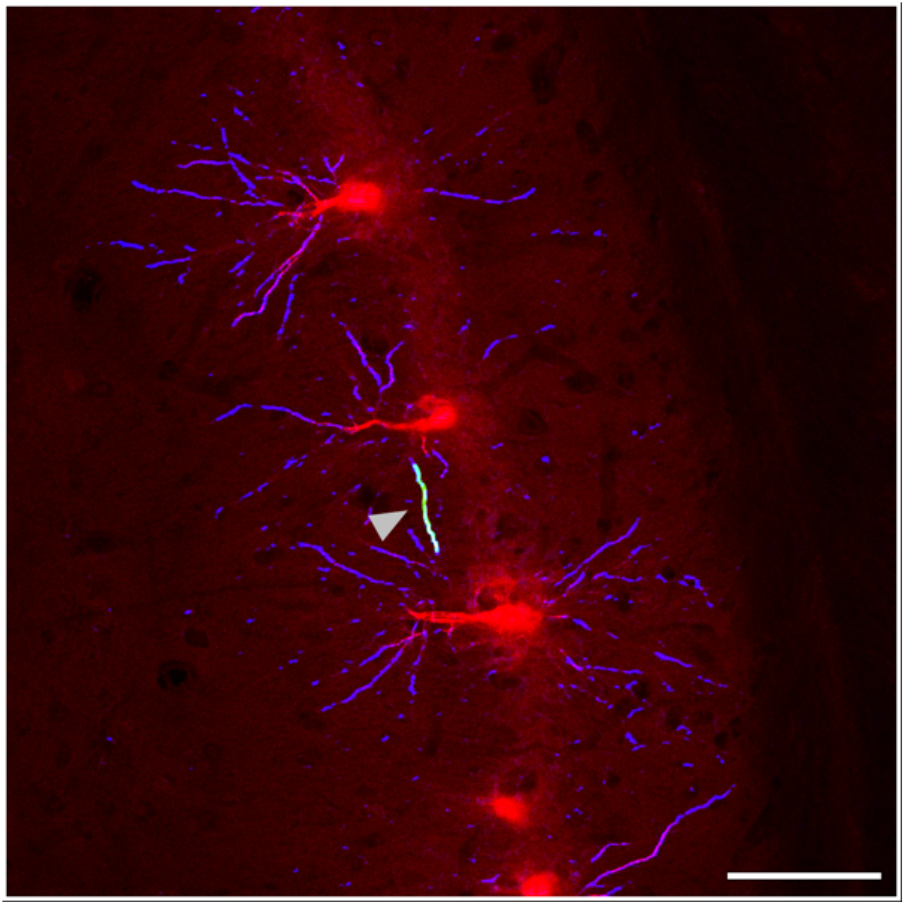
NAPA prediction of neurites in 3D coronal sections from fixed mouse brains. Fluorescence image in red, prediction in blue, provided annotation (marked with arrow) in green (appears white because of colocalization with prediction). Scale bar is 100 μm.

To distinguish between model quality for each tracing task in dataset III, and also more generally for datasets with available annotation, we used the black and white pixels in the manual traces to assign parts of an image to background (black) or foreground (white). The annotated foreground class accounted for only a few percent of the total pixels in the presented datasets, making it more difficult to learn. The background pixels were all non-neurite pixels, and a high background score represents the ability to remove soma, debris, and any other undesirable intensity sources. The foreground pixels were neurite pixels, and a high foreground score captures the model’s ability to correctly assign neurites within the annotated regions. A close up of a cell is shown in Figure 7 to illustrate the parts of the accuracy calculation. There are two caveats in this accuracy estimate: (1) many more background than foreground pixels are present in the image, so it is generally easier for background accuracy to be high, and (2) even the most thoroughly annotated images miss neurite intensities to different degrees, so some of the predicted intensities do get scored in the background class, negatively. This latter caveat makes a relatively small impact on the overall accuracy because neurite pixels make up only a few percent of the full image. From curation to accuracy evaluation, we have since applied this limited-curation strategy to new datasets, and can quickly obtain new models that predict neurite curation for new images.

**Figure 7.**
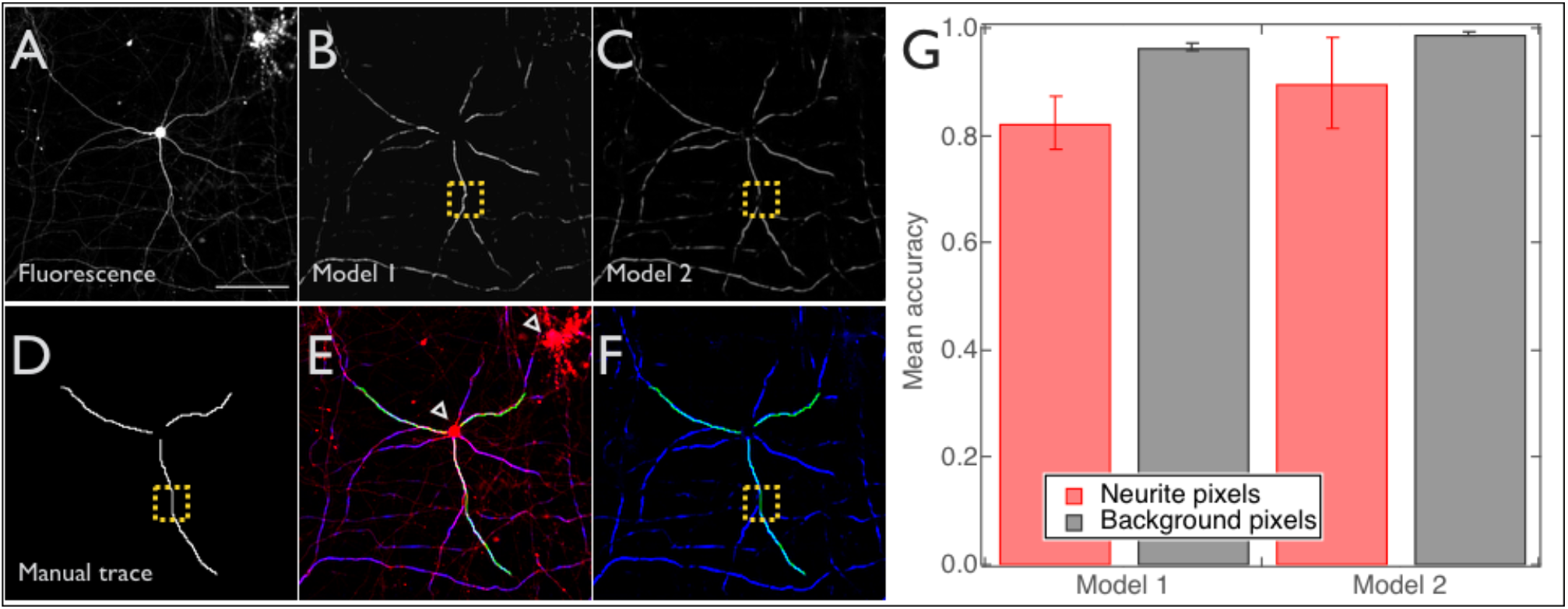
Model accuracy is evaluated based on the ability to capture ground truth. A. Fluorescence image of morphology marker. B-C. Examples of predictions made by two different sample models (different hyperparameter choices) for a single image. D. Annotated ground truth mask. E. Overlay of fluorescence (red), prediction from model I (blue), and annotation (green). Open arrows show regions that are correctly removed in the output image (background pixels in plot). F. Overlay of annotation (green) and prediction from model 2 (blue). Good overlap between prediction (blue) and annotation (green) results in high accuracy. Regions shown in green are scored as foreground (neurite pixels). All other regions are scored as background. Yellow boxes in B, C, D, and F show one point of difference between predictions from model 1 (B) and model 2 (C) within the scoring region (D) marked also by green in F. Differences such as this lead to different model accuracies. G. Model accuracy shown for two models applied to the same data. Red bars (neurite pixels) report the average fractional overlap of the white pixels in the prediction (B and C) and the annotation (D) within the annotated regions. Grey bars (background pixels) report the average fractional overlap of black pixels in the prediction (B and C) and the annotation (D) outside of the annotated regions. Error bars show standard deviation, and n = 12, 19 (model 1, 2). Scale bar in A is 100 μm.

Using the metrics outlined above, we next applied our top-performing model to generate full image predictions and quantify neurites across different conditions. Here, we aimed to determine whether the output could resolve the visual differences between iPSC-derived neurons from control and Parkinson’s disease samples (dataset IV). We globally analyzed neurite coverage in control and Parkinson’s disease neurons across eleven wells (annotated for this task), and compared fractional coverage of neurite pixels (total neurite pixels / total image pixels) in the disease and control wells. This fraction is calculated for each image representing a well. The predicted neurites showed lower fractional coverage in the disease wells than the control, in agreement with previous studies ^8,31–33^, and echoing the manually traced result (**Figure *8***).

One difference between the manually traced neurites and the predicted ones is that traced neurites have a fixed width that is propagated throughout their length, while the prediction tries to reproduce the actual width along the length of the neurite (**Supplementary Figure *1***). This difference affects the absolute magnitude we obtain from the manual traces. It is difficult to choose a generally relevant width, but we estimated 3 pixels for the datasets presented in this work. Comparison of the absolute numbers of pixels associated with neurites in manual and predicted traces showed that both methods capture a difference between the control and disease samples (**Figure *8***). To assess the extent of the difference, we compared the algorithm with multiple curators performing the same task (**Supplementary Figure *2***), and found that the neurite quantification from the model’s output detected a difference between the control and disease condition that was similar to the curators’ traces. The variation across curators is not a surprise, particularly since small differences in the display parameters has a noticeable impact on the visual curation landscape (example shown in **Supplementary Figure *3***). There are also notable differences between how the algorithm and curator perform the annotation task. For example, using an aided tracing tool we see that the manual traces sometimes take a shorter path in connecting neurites (**Supplementary Figure *4***). We observed that in addition to predicting the full and changing width of the neurite (**Supplementary Figure *1***), the model predicted many more of the intensities representing short neurites (Figure 8), and often followed the neurite more accurately than manual tracing. By contrast, manual curation tended to focus on the longer traces (Figure 8), meaning the frequent but shorter neurites pixels are missed. This could lead to a larger overall difference between disease and control conditions, an element that can be alleviated by an algorithmic approach.

Neurites are dynamic structures, and vital information relevant to health and disease could be gleaned by studying how neurites change over time. Unfortunately, because quantifying neurites manually at even a single time point is so laborious, quantifying how neurites change dynamically over time at scale is often unfeasible. To determine whether our algorithms could work for this purpose, we next looked to see how neurite arborization evolved over time from initial plating in the same eleven wells that were manually curated in **Figure *8***. **Figure *9*** shows the progression of neurite morphology over eight days. We found that the fractional neurite coverage began to differ between the disease and control samples after about 36 hours. In the control samples neurite arborization proliferated, while in the disease samples the neurite coverage stayed roughly constant and perhaps slightly dropped, which may represent loss or retraction of neurites commonly seen in neurodegenerative disease ^8,31^. The manually annotated time point in Figure *8* is the 4th point in **Figure *9*** plot (1.5 days). Extending this analysis to the rest of the time points required running the raw images through the DL model, but did not require extra time spent manually annotating each image, allowing us to further scale these types of quantifications to new conditions and questions ahead. The highly significant differences underscore the value of our automated approach for quantifying a potentially disease-relevant neurite phenotype that would otherwise be difficult or unfeasible to quantify manually.

**Figure 9.**
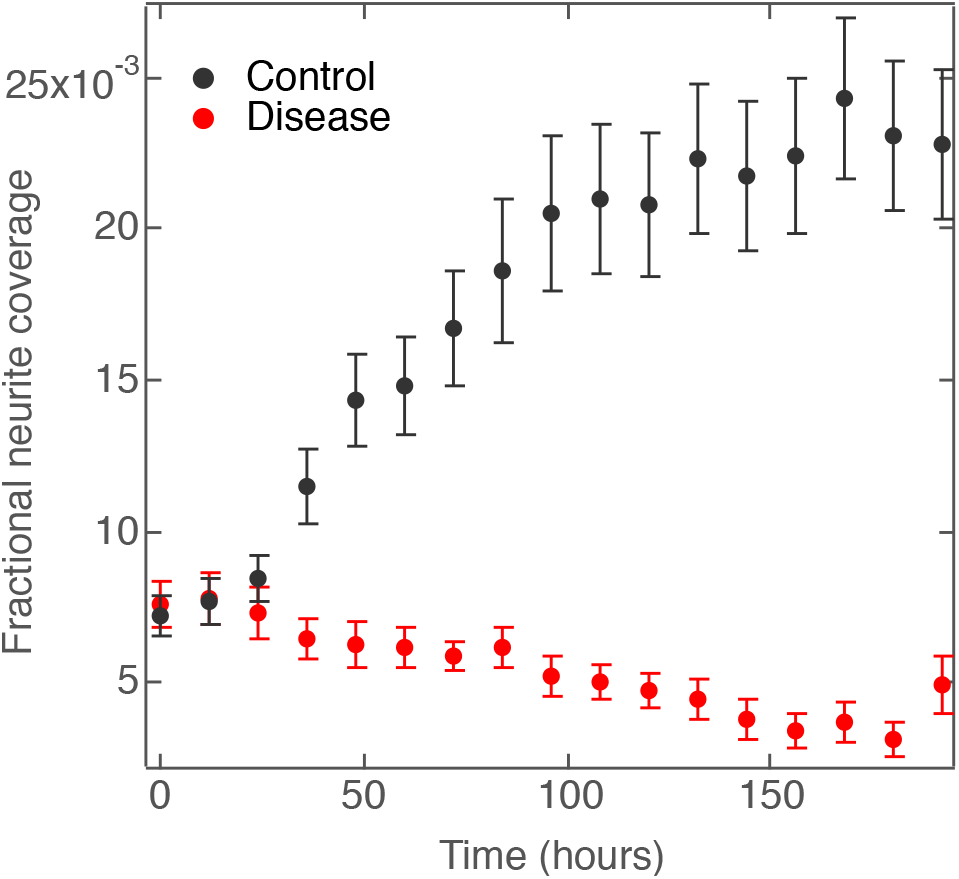
Longitudinal analysis of neurites in iPSC-neurons from control and Parkinson’s disease patients. For neurites in the healthy control cells, fractional neurite coverage increased over time, as expected, with cell proliferation. On the other hand, in cells from a Parkinson’s disease line, fractional neurite coverage stays low. The fourth time point corresponds to the images in Figure 8.

## Methods

### Datasets

All datasets were prepared as part of projects unrelated to the machine learning task described here. The culture and imaging conditions are described below. In this work, only the morphology marker was used, even though cells were transfected with multiple plasmids and fluorescence markers. For neurite annotation, only dataset III was traced specifically for the machine learning task.

### Culture for dataset I (Dopaminergic neurons)

iPSCs were grown on Matrigel (Corning)-coated plates and fed with mTeSR media (StemCell Technologies). iPSCs were differentiated into dopaminergic neurons as previously described ^34^. Cells were dissociated using trypsin-EDTA (ThermoFisher Scientific) and plated onto 384-well plates coated with poly-L-ornithine (Sigma-Aldrich), fibronectin (Corning), and laminin (Sigma-Aldrich). Biosensor expression and imaging: iPSC-derived dopaminergic neurons were virally transduced with AAV1-hSyn-GFP (Addgene). Cells were imaged at day 46 of differentiation and neurites traced using the NeuronJ plugin for ImageJ.

### Culture and imaging for dataset II (Primary rodent neurons)

Rat cortices were dissected from embryonic day 18–21 rats, digested with papain (10 U/ml) and seeded at 0.6 × 106 cells/ml on a poly-lysine/ laminin substrate in a 96 well plate (PerkinElmer #6005558). Primary cortical neurons were cultured for 5 days and then transfected using Lipofectamine 2000 (ThermoFisher # 11668019). Cells were incubated with 0.5 μl Lipofectamine 2000 and 100ng pGW1-mApple plasmid construct per well as a morphology marker as described^35^. Cells were incubated for no more than 20 min at 37° C before rinsing. The remainder of the transfection protocol was per the manufacturer’s suggestions, resulting in an overall 1–5% transfection efficiency, which sparsely labels the cultures and facilitates single-cell microscopy studies.

### Histology for dataset III (CA1 pyramidal neurons)

To reconstruct the dendritic tree architecture of single CA1 pyramidal cells, TgDyrk1A mice were perfused with 4% PFA, and 150 μm coronal sections from the dorsal hippocampal CA1 region (Bregma, antero-posterior = −1.06 to −2.54 mm; medio-lateral 0.5 to 1.6 mm) were obtained with a vibratome. Intracellular injections were performed in CA1 pyramidal neurons by continuous current of fluorescent Lucifer Yellow (LY) as described (Elston, 1997; Elston, 2001). Briefly, pyramidal neurons located randomly throughout the dorsal CA1 area (Bregma, antero-posterior = −1.06 to −2.54 mm; medio-lateral 0.5 to 1.6 mm; Paxinos and Franklin, 2001) were selected for LY injection and further reconstruction (number of neurons/animals: wild type = 18/4; TgDyrk1A = 19/4). Images of single neuron apical trees for neuronal reconstruction were acquired with a confocal microscope (SP5 Upright; Leica Microsystems) with a 20x air objective (HCX PL APO CS 20.0×0.70 dry UV). A line scan of 1024 × 1024 pixels, 0.347μm wide z steps and 3-line intensity averages were used for imaging whole dendritic trees.

### Culture for dataset IV (Forebrain neurons)

iPSCs were grown on Matrigel (Corning)-coated plates and fed with mTeSR media (StemCell Technologies). iPSCs were differentiated into forebrain neurons as previously described ^36^. Cells were dissociated using trypsin-EDTA (ThermoFisher Scientific) and plated onto 96-well plates coated with poly-L-ornithine (Sigma-Aldrich), fibronectin (Corning), and laminin (Sigma-Aldrich). Biosensor expression and imaging: iPSC-derived forebrain neurons were transfected with 0.1 μg hSyn-mApple plasmid per well of a 96 well plate using Lipofectamine 3000 reagent (ThermoFisher Scientific). Forebrain neurons were imaged at day 40 of differentiation, and neurites traced using the NeuronJ plugin for ImageJ.

### Microscopy

Datasets I, II, and IV were imaged longitudinally using a Nikon Plan Fluor ELWD 20X/0.45 NA objective mounted on a Nikon Ti-E inverted microscope controlled by a custom plugin for Micro-Manager written in Java. Fluorescent excitation and emission spectra were generated with a Sutter Lambda XL Xenon arc lamp and full-multiband filter set (DAPI/FITC/TRITC/Cy5; excitation 560/25 and emission 607/36 for mApple fluorescence; and excitation 485/20 and emission 525/30 for GFP fluorescence). An Andor Zyla4.2 sCMOS camera with 2048 × 2048 pixels was used to capture images. Each well was imaged with 50ms (dataset I), 400ms (dataset II), or 250ms (dataset IV) exposure, as 16 (4×4) or 9 (3×3) fields of view and images were background corrected and then stitched together using custom image processing scripts. The morphology images were used towards neurite tracing and model input for neurite prediction.

### Neurite tracing in FIJI without NeuronJ (used for dataset III)

Neurites were traced manually in FIJI using the NeuronJ plugin ^14^ with the freehand selection tool. Each trace was added to the ROI manager. After all neurites were traced, a new mask (all zeros) of the same dimensions as the original traced image was created. The collected ROIs were applied to the mask with pixel width of 3. The ROIs were flattened and mask image was binarized such that all neurite-traced pixels had values of 255 and background pixels were all zero. This binary mask served as paired input with the original image.

### Neurite tracing aided with NeuronJ (used for datasets I and II, and IV)

Alternatively, images were first converted to 8-bit RGB tiff images and then opened using the NeuronJ plugin. Brightness and contrast were first adjusted to enable visualization of neurites, and pixel width was adjusted to 3. Neurites were then traced starting at the cell body and following neurites outward. A snapshot that included traces was captured and converted to a binary mask, such that all neurite-traced pixels had values of 255 and background pixels were all zero. These binary images were paired with original raw images and comprised model inputs. Since subsequent steps begin with binary masks of traces, the two methods we used for manual neurite tracing were indistinguishable.

### Image processing

Images for each well were collected, background corrected, and stitched together into a montage using custom scripts.

### Model

We used several datasets to train a fully convolutional model that aims to predict neuronal morphology curation that reproduces the gold standard of manual curation. We used multiple datasets, with varying imaging and experimental conditions (Table 1). Each dataset was prepared to generate paired inputs for the model consisting of raw 16-bit images and curated masks of traced neurites (**Figure *3***). Each input pair was a 128 by 128-pixel square that came from tiling the original image and the target image (a neurite mask) at 128 pixel intervals. In our datasets, one image thus produces 1000 to 2000 tiles. The raw image input went through a sequence of convolution and max pooling steps for the model to learn kernel weights that minimize root mean squared error (RMSE) between predicted output and the curated mask. The curated mask was a black and white binary image where all background pixels were black and traced neurite pixels were white. Each tile was tracked with a label that made retrieval easy to reference to the original dataset and corresponding curation. Normalization was performed on the complete dataset to generate 128 by 128 normalization tiles used to scale each input tile from −1 to 1, and centered around 0. During normalization, we also calculated the total number of pixels belonging to neurites according to the curation. This number tended to be less than 2%, even though in training, we only used raw image tiles that have annotation (whose target/mask had white pixels). We trained only on data that had annotation (however little) to focus resources on learning the signal. Because neurite pixels (<2%) were so much rarer than the background pixels (98%), we used the fraction of neurite pixels to weight the calculated loss. Without loss weighting, the loss decreases quickly and the model learns to predict all pixels as background without much penalty.

Once tiles were generated, they were augmented with typical flips, rotations, and reflections available in the Tensorflow library, and the model was trained on this augmented input. We used tf.Dataset to further repeat, shuffle, and batch the input. The autoencoder consisted of an encoding/contracting path and a decoding/expanding path. The contracting path was a series of three convolution and max pool layers with ReLU activation. The expanding path consisted of a symmetric set of three convolutions and up-pool layers with ReLU activation. The complete model is shown in **Figure *3***a. Loss was defined as the mean square difference between predicted and actual pixel intensity for every pixel. The squared difference was multiplied by the weighted label vector which accounts for the low occurrence of neurite pixels. The weighting depended on the fraction of neurite pixels calculated for the full training dataset. Then, each non-neurite pixel was scored with an RMSE and multiplied by a factor that makes the contribution from non-neurite and neurite pixels about equal (rather than <2%). The RMSE loss was minimized to obtain the optimal weights. Approximately 1200 training steps reached loss-convergence with RMSE values ranging roughly 0.0002 to 0.002 for similar quality models.

We tested the effect of varying hyper-parameters on model quality. These parameters included training batch (50-500), kernel (3-11), number of layers (2-4), and whether or not weighting is used. For higher signal tasks, such as soma segmentation, 2 sets of convolutions and max pooling layers on the contracting path and 2 more sets on the expanding path were sufficient. All of the neurite work is lower signal and noisier, and has so far required 3 contracting convolutions and max pooling pairs, and 3 expanding convolutions and up-pooling. The higher batch size helps prevent over fitting which otherwise occurs early in training. Weighting of the loss focused the model to learn non-black pixels, which comprised a small fraction of the total pixels, typically close to 2%. Kernel size was affected the most by image properties. We saw that lower signal and noisier images performed better with larger kernel choice. The models whose predictions are shown here all used a kernel of 3 pixels.

For all trained models, the datasets (128×128 image pairs) were split into training (90%), validation (5%), and holdout (5%) sets. The training tiles were used for model training. Each model was evaluated by RMSE score and also qualitatively for its ability to capture neuronal morphology outside of the annotated pixels –these are neurites that are visible in the raw image, but were not traced and are thus not part of the mask, which provided the ground truth for scoring. The visual evaluation was applied to the unseen-by-the-model testing dataset and the best models were selected based on these two criteria. The top models were then applied to the holdout dataset and their quality was verified by RMSE and by their ability to predict neurite annotation as well as remove soma and debris from the image (ability to capture neuronal morphology within the raw image). The images shown in all the figures reflect each model’s performance on the holdout data or different datasets, all unseen by the model during training or tuning.

### Accuracy metrics

We evaluated accuracy in several ways. The model training and evaluation of training used a mean pixelwise root-mean-square-error. Predicted pixels were compared to the target image (ground truth annotation) and mean squared error calculated. For each model, this metric guided improvement and hyper-parameter choice as the loss function. The training used only annotated patches of image, however, to evaluate model’s ability to reproduce the fully reconstructed annotation, we used accuracy within the ground truth regions of the whole image – all the annotated neurite pixels. Binary predicted images were compared to ground truth binary masks within the manually annotated neurite regions. The fraction of total neurite pixels that were also predicted as neurite pixels by the model was calculated per image. A mean and standard deviation across all images is reported in the plots. The mean fraction of total background pixels (soma, debris, all non-neurite pixels) that are also predicted as background pixels are shown side-by-side with the neurite pixels (**Figure *7***). To compare across conditions and quantify the number of neurites in an image, total neurite pixels were counted for each binary manual or predicted image and reported as a fraction of all pixels in the image. Error bars show variation across images within a condition.

In the case of the partially annotated datasets that we used to generate a model, the annotated dataset was divided as above into training, validation, and holdout sets. Once a model was generated and full images for the complete dataset were reconstructed, the initial annotated tiles were used within the dataset at prediction time, meaning that 450 tiles that were used in training were also part of the neurite prediction at run time where 3–8 million inputs required prediction.

## Discussion

Here we report a new approach to obtaining neurite traces from fluorescence images of neurons labeled with a morphology marker. We use a fully convolutional DL model that can selectively keep neurites and remove soma, debris, and other imaging artifacts that are common in high throughput and longitudinal imaging.

Neurite annotation is challenging due to several factors. First, the dimness of the neurites creates inherent signal to noise constraints. The neurites change in intensity throughout their length and, at such low signal, the background is uneven as well, especially in non-confocal, epifluorescence imaging. Further, the background near larger, thicker, and highly fluorescent objects (soma, cellular debris, or other artifacts) is often significantly greater than near dim objects. Thus, neurite tracing often requires multiple zoom levels and contrast updates to accurately assign the source of signal individually and contextually. Due to these complexities, it is also difficult to simplify the tracing task with conventional image processing steps, which typically begin with selecting a single pixel intensity to demarcate foreground from background. The choice of intensity at this early step presents an inherent trade-off. Either the threshold is set high, and the dim thin neurites fall below the level of detection, or the threshold is set low, and the portions of the background adjacent to bright objects (e.g., the thickest part of neurites, near the soma) are assigned to foreground, also leading to erroneous measurements.

Previous approaches to automate this process broadly fall into two categories: tools that help annotate, and tools that take parameters from an image to search for neurite-like patterns. The tools that trace neurites (rather than assist in manual annotation), are often based on pipelines that combine image pre-processing, neurite tracing, and subsequent post-processing compatible with downstream quantification ^37^. Some solutions include novel ways of tracing out a neuronal projection from a seed soma ^38^, human in the loop ^39^, computer generated input ^40,41^, or using combinations of filters/transformations to skeletonize images and extract neurites ^42–44^. Other methods follow gradients ^44,45^ and minimize an energy function. Some methods are substantially more computationally intensive than others ^16^, and some require a lot of tuning parameters ^14^, but if they generalize, are much faster. However, in high throughput or longitudinal imaging contexts, where the experimental conditions can change dramatically over the course of the imaging experiment, parameter generalization is a particularly challenging task. DL-based algorithms can help capture relevant parameters more broadly and overcome experimental variation inherent to fluorescence imaging.

DL approaches have previously been applied in the assignment of foreground and background pixels to perform various segmentation tasks ^30,46^. In fact, several deep learning approaches ^38,41,47–49^ have been applied to neurite tracing, a related task. Our DL network is based on an autoencoder architecture ^50–54^ with a fully convolutional approach, where our task is instead to de-noise the images by removing any non-neurite intensities. With a related, U-Net architecture applied to the same datasets, we saw segmentation output often returned soma in addition to neurites, thus failing to classify soma pixels as background. We showed that using the lightweight autoencoder-based architecture, it is possible to predict neurites across multiple datasets. We also analysed diverse sample types to better understand the breadth of neuronal data variation. This allowed us to come up with a compact scheme for generating predictive models from a limited number of annotated inputs, an annotation task that should be possible to achieve in less than an hour. Our approach enables quick training towards new datasets. Because a single dataset might be comprised of thousands of images, the relative cost of generating a custom model is still low, as the model enables generation of a fully curated dataset from relatively few curated examples (about 30– 40 traced neurites per dataset). Further, we verified that this approach removes undesirable intensities and correctly returns neurites at about 80% accuracy compared with manual annotation, which unfortunately is itself error-prone ^11^. When we applied NAPA to a larger dataset from a 96-well neuronal culture of healthy control and Parkinson’s disease lines imaged over 192 hours we were able to resolve differences between these two conditions and quantify neurite outgrowth over the time course of the study, a task that would be quite difficult to achieve with manual annotation alone.

Indeed, the predictions made by NAPA are not perfect. Neurites are still missed, particularly in very neurite-dense cultures, and those that are predicted are often not completely continuous, which is problematic for per-cell quantifications downstream. Indeed, potential per-cell quantifications are generally complicated by ownership ambiguity in denser cultures, even with continuous predictions. However, an advantage with our approach is that the prediction output reflects the annotation in the training data, which provides a direct readout for what types of events are poorly represented in the training data. This can inform a more narrowly targeted curation to not only add examples of what the NAPA has missed and improve its models, but also to address new needs. Since NAPA amplifies the training data, the aim of curation is to teach the models what the curator would like traced the image, which can be tailored to the task. Output images could also be generated for the purpose of broadly improving signal to noise ratio in the fluorescence images ^55^ and interface with post processing algorithms that focus on reconstruction of island neurite intensities (not presented) as has been achieved with other algorithms ^47,56^.

Since the annotation we used was not directly generated for the DL task, its coverage varied, depending on the sample and experimental conditions. Very dense networks of neurites, covering more than 10% of the pixels in an image (> 1 million neurite pixels in the examples here), did not have every neurite annotated; more complete annotation coverage is reasonable for sparser samples, although it is still difficult and requires multiple iterations with different contrast settings. The main concern with partially annotated data was that false negatives would be introduced in training. We initially tried minimize false negatives by controlling the context provided with the size of the input patch. However, we soon learned that by weighting the loss towards the annotated pixels, the model was able to focus on correctly predicting the various types of positive neurite examples provided in the training data, and in effect amplify them in the rest of the image. This technique enabled us to ask more general questions about neuronal morphology.

There is certainly a trade-off between manual tracing and automated techniques. On one hand, manual tracing may be highly accurate, although not perfect in its ability to faithfully follow neurites (**Supplementary Figure *4***) or in its completeness. The problem is that the manual traces remain bound to the dataset and do not extend to very large datasets, including longitudinal and dynamic ones, where tracing is not a feasible option. In addition, the traces generated are prone to bias based from the curator, and as we have seen, the same dataset can be traced to yield different outcomes (**Supplementary Figure *2***). On the other hand, fully-automated, off-the-shelf algorithms have significant limitations, requiring substantial manual tuning or exhibiting brittle performance that is susceptible to the qualities of the dataset. Another aspect of this trade-off is time. This can be time allocated to manual tracing or time needed to tune parameters to a given dataset. In the case of manual annotation, to fully annotate an image with culture density shown in **Figure *8*** would require 20–40 minutes per image using an assisted tracing tool such as NeuronJ. Denser samples require more time. With 30 minutes per image, a typical dataset of 96 wells and 9 timepoints would requires 432 hours to manually annotate. Further, the variation between curators (**Supplementary Figure *2***) might increase the required time. With our DL approach, the per image curation time is closer to 10 minutes. We typically annotate about 20 representative images from the entire experiment, so this would amount to ~3 hours of annotation in the slowest case (without tracing assistance from NeuronJ), and a 100x speedup. For a purely manual curation approach, as the experiment grows, the manual curation time needed would scale and likewise grow. With the approach we describe, the manual curation time would remain constant. The strategy and algorithm described here are an effort to bring together the best parts of manual and automated annotation by using an ML algorithm that requires little input.

Neuronal morphology plays an important role in signaling, development, and disease, but its study is limited by the labor-intensive process of manually generating cell traces. Although manual tracing is technically possible for any task, it can be so impractical and expensive for larger datasets that it is effectively unfeasible. Hence, automating neuronal curation would enable us to address broader questions more quantitatively and with improved statistical power. Here we present NAPA, a CNN-based DL approach to predict manual curation from a small fraction of traced neurites, and show quantification applications that allow us to broadly distinguish between disease and control samples and over time. In each case, the input to the model is the fluorescence image and the annotation to be amplified, and the output is ultimately a binary mask that can be used with already available algorithms. Since this supervised model learns from given examples, it can be targeted to more specific questions in the initial annotation step, such as annotating branch points or a particular type of neurite break or contact. These kinds of neurite features would be difficult to access through current technologies that focus automation on more general neurite properties.

## Acknowledgements

We thank all the members of the Finkbeiner and Dierssen laboratories for helpful discussions and advice throughout the preparation of the manuscript. We thank Guangzhi Li and Vivek Ramaswamy for implementing several algorithms from the literature for our evaluation, Sara Modan and Preeti Rao for tracing of the 11 wells for the PD vs control i-neurons, Margarita Fedorova for dataset management and processing, Stephen Cannon for initial steps towards neurite ownership (not presented here). We are grateful to Sarah Gardner for graphical design of Figure 1, and Kathryn Claiborn for editorial assistance. This work was made possible with support from the Koret Foundation Artificial Intelligence Program for Biomedical Research at Gladstone, and NIH grants RF1 AG058476, P01 AG054407, R37 NS101996 and U01 MH115747. The lab of MD is supported by the Generalitat de Catalunya (2017 SGR926). We acknowledge the support of PID2019-110755RB-I00/AEI / 10.13039/501100011033, Horizon 2020 grant agreement No 848077, NIH 1R01EB 028159-01, JPND Heroes and JPND-NIH. We acknowledge support of the Spanish Ministry of Science and Innovation to the EMBL partnership, the Centro de Excelencia Severo Ochoa and the CERCA Programme / Generalitat de Catalunya. The CIBER of Rare Diseases (CIBERER) and, CIBER of Physiopathology of Obesity and Nutrition (CIBEROBN) are initiatives of the ISCIII.

## Footnotes

5 The minimum and maximum intensity values in an image capturing the dynamic range. All images are 16-bit with possible values of 0 to 65,53565535.

6 The mean intensity standard deviation for the image.

7 Annotation coverage crudely captures how many of the imaged neurites are traced. Complete: more than 80% of the neurites are traced for model training. Partial: three of the most prominent neurites were traced for several cells in the image for model training. None/partial: No neurites were annotated for model training and some annotation was added only for the purpose of comparison and evaluation. For dataset III, the partial annotation was later used for training a new model.

8 Dopaminergic neurons

9 CA1 pyramidal neurons from TgDyrk1A Down syndrome mouse model.

## Notes

### Competing Interest Statement

The authors have declared no competing interest.

