## Supplemental information for "A deep learning approach to neurite prediction in high throughput fluorescence imaging"

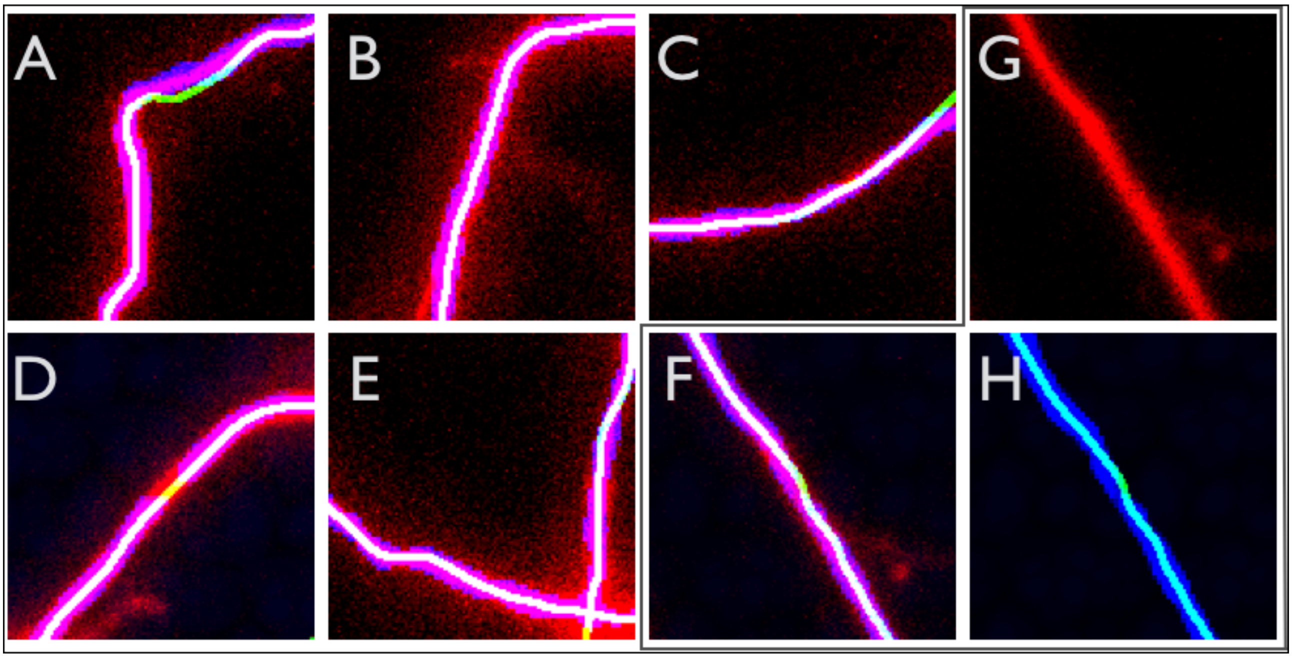


**Supplementary Figure 1.** Differences in trace vs algorithm. Manual trace has a uniform width of a set number of pixels (shown here, 3), while the algorithm tries to capture the full width of the neurite and how it changes throughout its length. A–F. Close up examples of neurites showing overlays between fluorescence image (red), manual trace (green), and prediction (blue). Good overlap agreement yields a white color. G–H fluorescence intensities (G) and prediction and manual trace (H) show the contributions of each channel more clearly for example F. All panels have the same scale and are 40 μm per side.

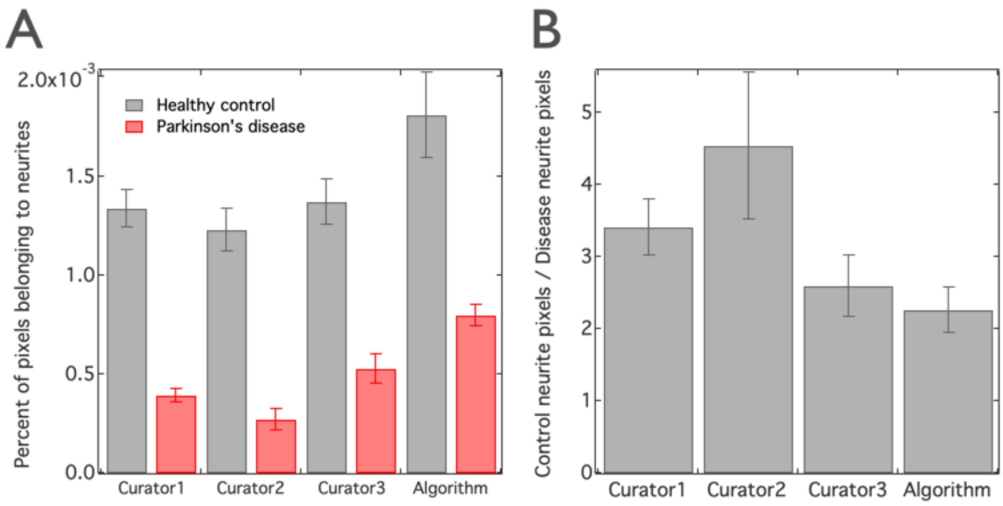


**Supplementary Figure 2.** Variation across three curators performing the same tracing task on 11 images as in **Figure 8** and **Figure 9**. Despite using the same tracing assist program and performing manual tracing on the same 11 wells, there was some curator-to-curator variation in neurite quantification. Curator 2's trace shows the largest difference between control (grey) and disease (red) samples. Curator 2's annotation is used in the evaluation in **Figure 8** and **Figure 9**. The ratio of control and disease for each curator is shown in the right panel. Error bars show standard error. Control bars represent 6 images and disease bars represent 5 images. The ratio representing the fold higher fractional coverage of neurites for control over disease samples falls within error across curators and algorithm.

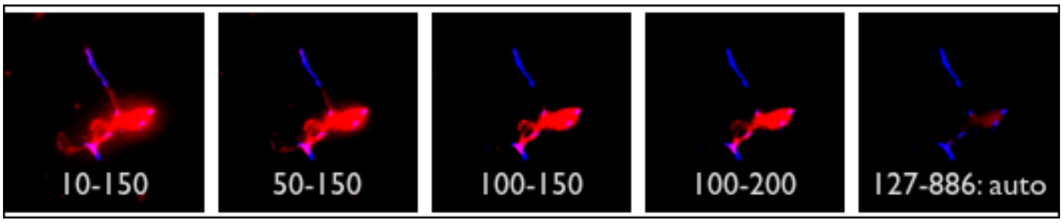


**Supplementary Figure 3.** An example of contrast variation showing a dimmer cell (red) that does not receive manual trace, with prediction (blue) shown across different contrast settings for the fluorescence channel (red). The numbers show minimum and maximum intensity range settings for contrast. The right-most image is the default auto-contrast setting for the full image based on the range of intensities in the rest of the sample. This sequence of images shows the effect of contrast settings on the curator's visual tracing landscape. Depending on the contrast parameters, some neurites are not visible and are less likely to be curated.

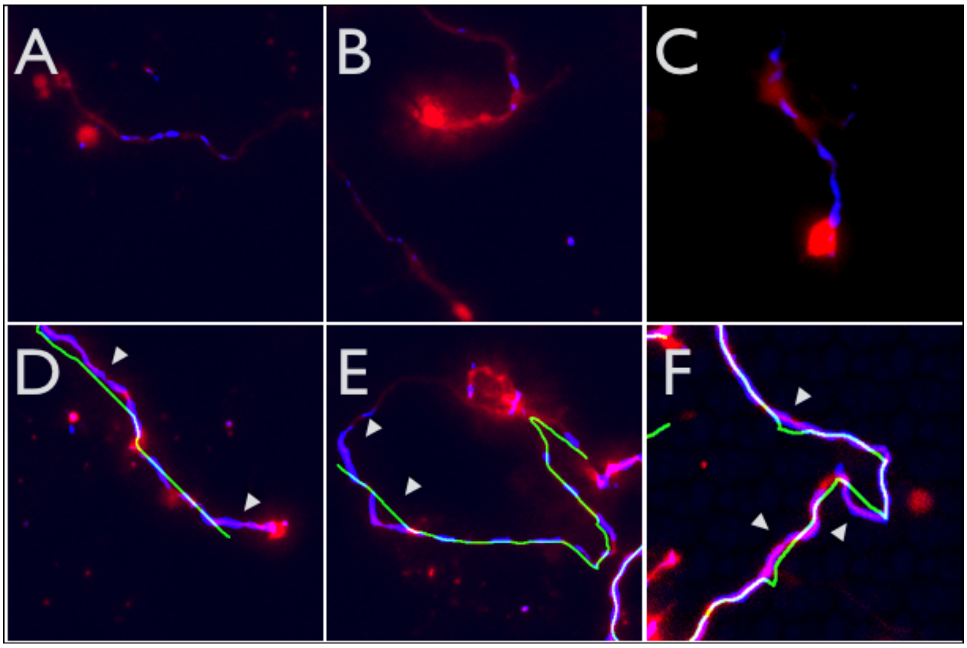


**Supplementary Figure 4.** Examples of curation variation. Close up regions of neurites showing differences between the manual trace and the algorithm's neurite prediction. A–C. Examples of neurite predictions (blue) in regions where no manual trace (green) was provided. These tend to be much dimmer intensities, and contrast was adjusted to saturate fluorophores (red) for visual clarity. D–F. Common longer-range variation between algorithm (blue) and manual trace (green). These differences result from the trace-aiding program used to manually generate ground truth (green). Grey arrows indicate regions of disagreement between manual trace, neurite intensity, and prediction. We also see examples of regions where neurite intensities have no associated manual trace and no associated prediction. Unlike the manual trace, model predictions typically do not appear in regions without cell intensity, but sometimes do miss neurite pixels, based on the seen training data. All square panels are 166 x 166 μm.

**Supplementary Movie 1.** Neurite annotation through tissue in 3D. Application of NAPA to 3D coronal sections from fixed mouse brains. Movie shows different heights in the z-dimension (thickness). Fluorescence image in red, prediction in blue, provided annotation is in green (appears white because of colocalization with prediction). Scale bar is 100 μm.
